## Supplementary Information for "Energy-based Graph Convolutional Networks for Scoring Protein Docking Models"

| | |
|---|---|
| Training Set | 1N8O 7CEI 1DFJ 1AVX<br>1AY7 1IQD 1CGI 1EZU<br>1JPS 1R0R 2FD6 2I25<br>2B42 1EAW 2JEL 1PPE<br>1BJ1 1KXQ 1BVN 1EWY<br>1KAC 1OPH 1E6J 2HLE<br>1WEJ 1A2K 1RLB 1GLA<br>1E6E 1J2J 1BUH 1XQS<br>1M10 1IJK 2HRK 1GP2<br>1GRN 1IBR 1BKD 1Y64 |
| Validation Set | 1ATN 1ACB 1AHW 1AK4<br>1AKJ 1AZS 1B6C 1BVK<br>1XU1 4CPA |
| Benchmark Test Set | 1DQJ 1E4K 1E96 1EER<br>1EFN 1F34 1F51* 1F6M<br>1FAK 1FC2 1FCC* 1FFW<br>1FLE 1FQ1* 1FQJ 1FSK<br>1GCQ 1GHQ 1GL1 1GPW<br>1H9D 1HCF 1HE1* 1HIA<br>1I2M 1I4D 1I9R* 1IB1<br>1IRA* 1JIW 1JK9* 1JMO<br>1JTG 1JWH 1JZD* 1K4C*<br>1K5D 1K74* 1KLU 1KTZ<br>1KXP 1LFD 1MLC 1MQ8<br>1NCA 1NSN 1NW9 1OC0<br>1OFU* 1OYV* 1PVH 1PXV<br>1QA9 1QFW* 1R6Q 1R8S<br>1RV6 1S1Q 1SBB 1SYX*<br>1TMQ* 1UDI* 1US7 1VFB<br>1WQ1 1XD3 1Z0K 1Z5Y*<br>1ZHH* 1ZHI 1ZLI 2A5T*<br>2A9K 2ABZ 2AYO* 2B4J<br>2BTF 2CFH* 2G77* 2H7V*<br>2HQS 2I9B 2IDO* 2J0T<br>2J7P* 2MTA 2NZ8* 2O3B<br>2O8V* 2OOB 2OT3 2OUL<br>2OZA 2PCC 1CLV* 1D6R*<br>2SIC 2SNI 2UUY 2VIS<br>2Z0E* 3CPH 3D5S* 3SGQ* |

| | |
|---|---|
| | 1YVB 9QFW* BOYV* |
| CAPRI Test Set | 2REX* 2WPT 3BX1* 3FM8* 3Q87* 4G9S* 4JW2* 4JW3* 4OJK* 4QKO* 4UEM* 4UF5* 4UHP* 4XL5* |
| Score_set, CAPRI Benchmark Test Set | 2VDU* 2R20* 3BX1* 2W5F* 2W83* 3FM8* 3FM8* 3E8L* 2WPT 3Q87* 3U43* 4JW2* 4JW3* |

**Table S1**. Protein complexes (represented by their 4-letter PDB IDs) used in training, validation, and 3 test sets. The PDB IDs with stars correspond to cases where native binding affinities were predicted from protein sequences.

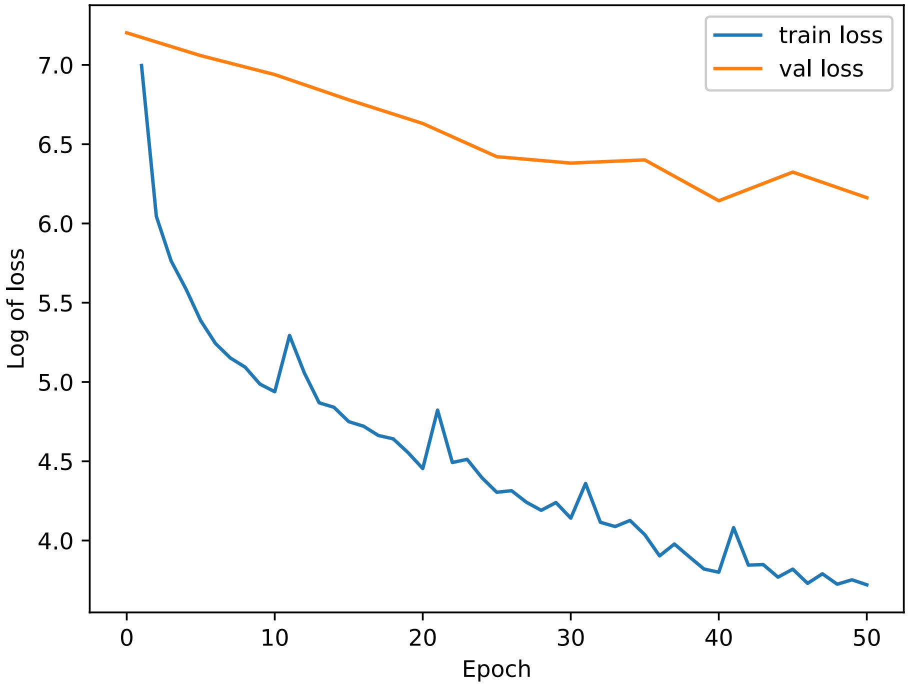


**Figure S1**. Training and validation loss over epochs during the training of the EGCN model.
